# Distinct structural transitions of chromatin topological domains coordinate hormone-induced gene regulation

**DOI:** 10.1101/003293

**Authors:** François Le Dily, Davide Baù, Andy Pohl, Guillermo Vicent, Daniel Soronellas, Giancarlo Castellano, François Serra, Roni H. G. Wright, Cecilia Ballare, Guillaume Filion, Marc A. Marti-Renom, Miguel Beato

## Abstract

The human genome is segmented into Topologically Associating Domains (TADs), but the role of this conserved organization during transient changes in gene expression is not known. Here we described the distribution of Progestin-induced chromatin modifications and changes in transcriptional activity over TADs in T47D breast cancer cells. Using ChIP-Seq, Hi-C and 3D modelling techniques, we found that the borders of the ∼2,000 TADs in these cells are largely maintained after hormone treatment but that some TADs operate as discrete regulatory units in which the majority of the genes are either transcriptionally activated or repressed upon hormone stimulus. The epigenetic signatures of the TADs are coordinately modified by hormone in correlation with the transcriptional changes. Hormone-induced changes in gene activity and chromatin remodeling are accompanied by differential structural changes for activated and repressed TADs. In response to hormone activated TADs exhibit higher density of internal contacts, while repressed TADs show less intra-TAD contacts. Integrative 3D modelling revealed that TADs structurally expanded if activated and compacted when repressed, and that this is accompanied by differential changes in their global accessibility. We thus propose that TADs function as “regulons” to enable spatially proximal genes to be coordinately transcribed in response to hormones.

## Introduction

The three-dimensional (3D) organization of the genome within the cell nucleus is non-random and might contribute to cell-specific gene expression. High-throughput Chromosome Conformation Capture (3C; (Dekker et al. 2002) derived methods have revealed that chromosome territories are organized in at least two chromatin compartments, one open and one closed, which tend to be spatially segregated depending on their transcriptional activity (Lieberman-Aiden et al. 2009). At a finer level of organization, some functionally related genes have been shown to be brought close in space to be transcribed in a correlated fashion during cell differentiation. These genes, which can be located on different chromosomes, are organized in spatial clusters and preferentially transcribed in the same “factories” (Osborne et al. 2004; Cavalli 2007; Osborne et al. 2007). Whether such mechanisms participate in transient modifications of the transcription rate in differentiated cells responding to external cues is still unclear (Fullwood et al. 2009; Kocanova et al. 2010; Hakim et al. 2011). Transient regulation of gene expression at the transcription level depends on the establishment of regulatory loops between regulatory enhancers and the promoter of responsive genes. Recent studies have demonstrated that these networks of interactions and the precise molecular mechanisms were more complex than initially thought (Shen et al. 2012; Thurman et al. 2012). Notably, a recent study showed that TNF responsive genes promoters are unexpectedly found juxtaposed to TNF responsive regulatory elements prior to stimulation, supporting the importance of the three-dimensional chromatin landscape in cell specific gene regulation (Jin et al. 2013).

Recently, an intermediate level of partitioning of the chromosomes in so called Topologically Associating Domains (TAD) has been revealed (Dixon et al. 2012; Hou et al. 2012; Nora et al. 2012; Sexton et al. 2012). Changes in the levels of expression of genes appeared to be correlated within TAD during cell differentiation (Nora et al. 2012), potentially by topologically constraining enhancer or silencer activity (Symmons et al. 2014). However, whether the spatial structure of TADs in differentiated cells plays a role in regulating the transient response to external signals in terminally differentiated cells remain mainly hypothetical (Nora et al. 2013). We have addressed this issue using T47D cells, a Progesterone Receptor (PR) positive breast cancer cell line (Truss et al. 1995) that respond to Progestins with transient changes in gene expression. We have studied the relationship between hormone-regulated changes in transcription, chromatin structure, epigenetic marks and the three-dimensional structure of TADs. We know that the progestin-activated-PR cross-talks with kinase signaling networks to modify chromatin and to regulate the transcription rate of thousands of target genes, either by activation of repression (Migliaccio et al. 1998; Vicent et al. 2006; Vicent et al. 2011; Wright et al. 2012). Using RNA-Seq, ChIP-Seq, Hi-C and three-dimensional (3D) modeling, we confirmed that the T47D genome is organized into about 2,000 TADs, whose borders are largely stable upon hormone treatment. Some TADs showed a general biased transcriptional response to Progestin (Pg), with topological segregation of Pg-induced stimulation and repression of gene expression. Moreover, hormone treatment induced large-scale modifications of TAD chromatin structure, which correlated with changes in spatial interactions within the responsive TADs. Integrative 3D modelling revealed that those responsive TADs structurally expanded if activated and compacted if repressed. Such structural changes are accompanied by differential changes in accessibility of the whole domain suggesting a direct link between TADs structure and activity. We thus propose that TADs function as “regulons” to enable spatially proximal genes to be coordinately transcribed in response to hormones.

## Results

### Characterization of TADs in T47D cells

To test whether the spatial organization of the genome into TADs is important for the response of T47D breast cancer cells to progestin (Pg), we generated Hi-C libraries of cells synchronized in G0/G1 and treated or not with Pg for 60 min. In order to minimize artefacts linked to the biased genomic distribution of restriction enzymes (Yaffe and Tanay 2011), we generated biological replicates of Pg treatment using independently *HindIII* and *NcoI* (giving in total 174,278,292 and 169,171,077 pairs of interacting fragments respectively for −Pg and +Pg cells, respectively; see Supplementary Table 1). Contact matrices at various scales showed that, despite the chromosomal rearrangements in T47D cell line (Supplementary Fig. 1A), the genome conserves the general principles of organization described for other cell types, notably the segmentation of chromosomes into Topologically Associating Domains (TADs; Fig. 1A). After correction for technical biases, the contact maps obtained with *HindIII* and *NcoI* yielded similar TAD borders (Fig. 1A). These datasets were therefore pooled to robustly define the position of TADs using a novel algorithm based on a change-point detection approach (see Supplementary Material). By testing different resolutions, we achieved the most robust identification of TAD boundaries at 100 Kb, at which we identified 2,031 TADs with a median size of ∼1 Mb (Supplementary Fig. 1B, C). These were organized as described in other cell types (Dixon et al. 2012; Hou et al. 2012; Nora et al. 2012; Sexton et al. 2012) with enrichment of genes and active epigenetic marks near their borders (Supplementary Fig. 2).

**Figure 1:**
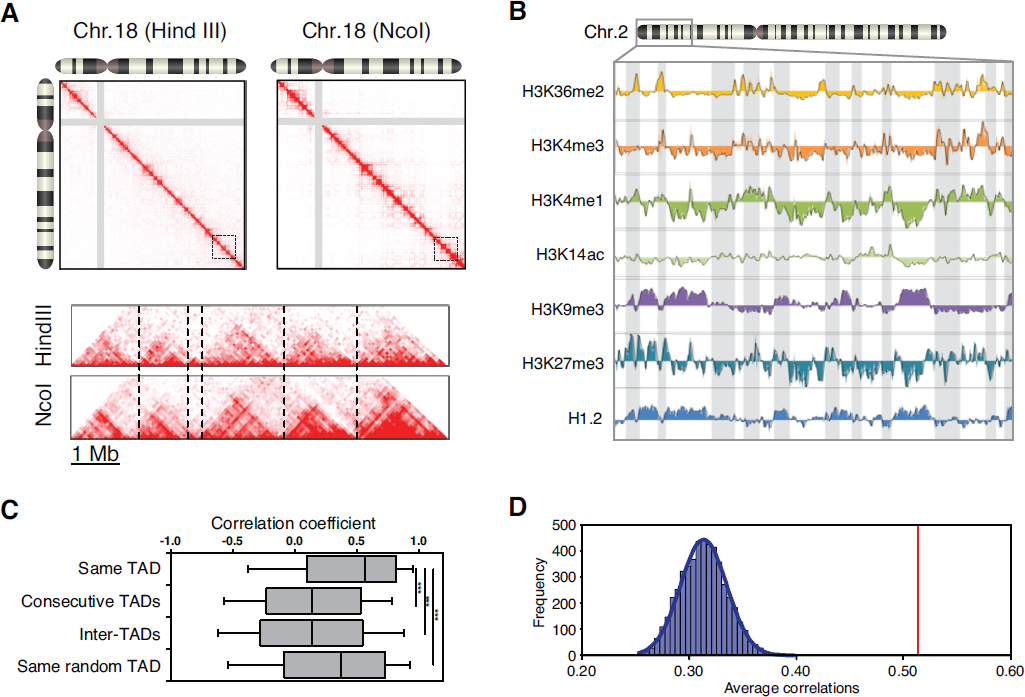
Topological chromatin landscapes defined in T47D cells. **(A)** Hi-C contact matrices of the chromosome 18 obtained using *HindIII* or *NcoI.* Both restriction enzymes resulted on similar detection of TADs borders (Bottom panel). **(B)** Log_2_ normalised ChIP-Seq/Input ratio on windows of 100Kb over a 30 Mb region of the Chromosome 2. **(C)** Distributions of pair-wise correlation coefficients of chromatin profiles between 100Kb chromosomal fractions located within the same TAD, within consecutive or randomly picked TADs or in a similar randomly defined domain for the region depicted in B. (***) *P* < 0.001; Mann-Whitney test. **(D)** Genome-wide analysis of the correlation of epigenetic marks between 100 Kb fractions belonging to a same TAD or to a same randomly positioned domain (see Supplementary Material). Histogram shows the distribution of genome-wide average correlation coefficients observed for 5,000 randomizations of TAD borders. The red bar (0.536) corresponds to the average correlation coefficients calculated for the real positions of the borders (*P* < 0.002).

To investigate the chromatin states of the identified TADs, we analysed various ChIP-seq experiments performed in this cell line under similar conditions of culture and treatment (Fig. 1B). We found that histone marks were frequently spread over entire TADs, which suggested that TAD borders delimit large chromatin blocks (Fig. 1B). Epigenetic marks were more homogeneous within than between TADs, and the transitions between high and low levels of the analyzed epigenetic marks occurred preferentially at the borders (Fig. 1B and Supplementary Fig. 3A) suggesting that TADs boundaries can act as barrier between epigenetic states. The correlation between combinations of epigenetic marks was higher within TADs than either between TADs or within random (either shifted or flipped) genomic domains of similar size (Fig. 1C, 1D). Together, these observations indicate that the TADs borders we defined in T47D are delimiting epigenetic domains and that transition between domains of different states preferentially occur at the boundaries between TADs, confirming previous studies (Dixon et al. 2012; Hou et al. 2012; Nora et al. 2012; Sexton et al. 2012).

### TADs respond as units to the hormone

We observed that around 90% of TAD borders detected without Pg treatment remained similarly positioned after Pg stimulation, with no evidence for specific, Pg-induced repositioning at this resolution (Fig. 2A). Thus, the T47D genome is organized in canonical TADs that do not significantly change boundaries in response to Pg. We previously showed that gene regulation by Pg involves a complex interaction of hormone receptors with kinase signaling pathways and chromatin remodeling enzymes to modulate the transcriptional output (Vicent et al. 2008; Wright et al. 2012). Here, we observed that the Pg-induced modifications of chromatin marks could occur over large genomic domains (Fig. 2B). Although these modifications of chromatin states could spread over consecutive TADs (Fig. 2B), transitions between divergent chromatin changes occurred more frequently at the TAD borders (Fig. 2B and Supplementary Fig. 3B). Importantly, the combinations of Pg-induced changes were more homogeneous within TADs than either between TADs or within random genomic domains of similar size (Fig. 2C, 2D), indicating that differential Pg-induced changes in chromatin marks levels are restrained within some TADs. Thus, TADs are epigenetic domains, the chromatin of which can be coordinately modified in response to external cues.

**Figure 2:**
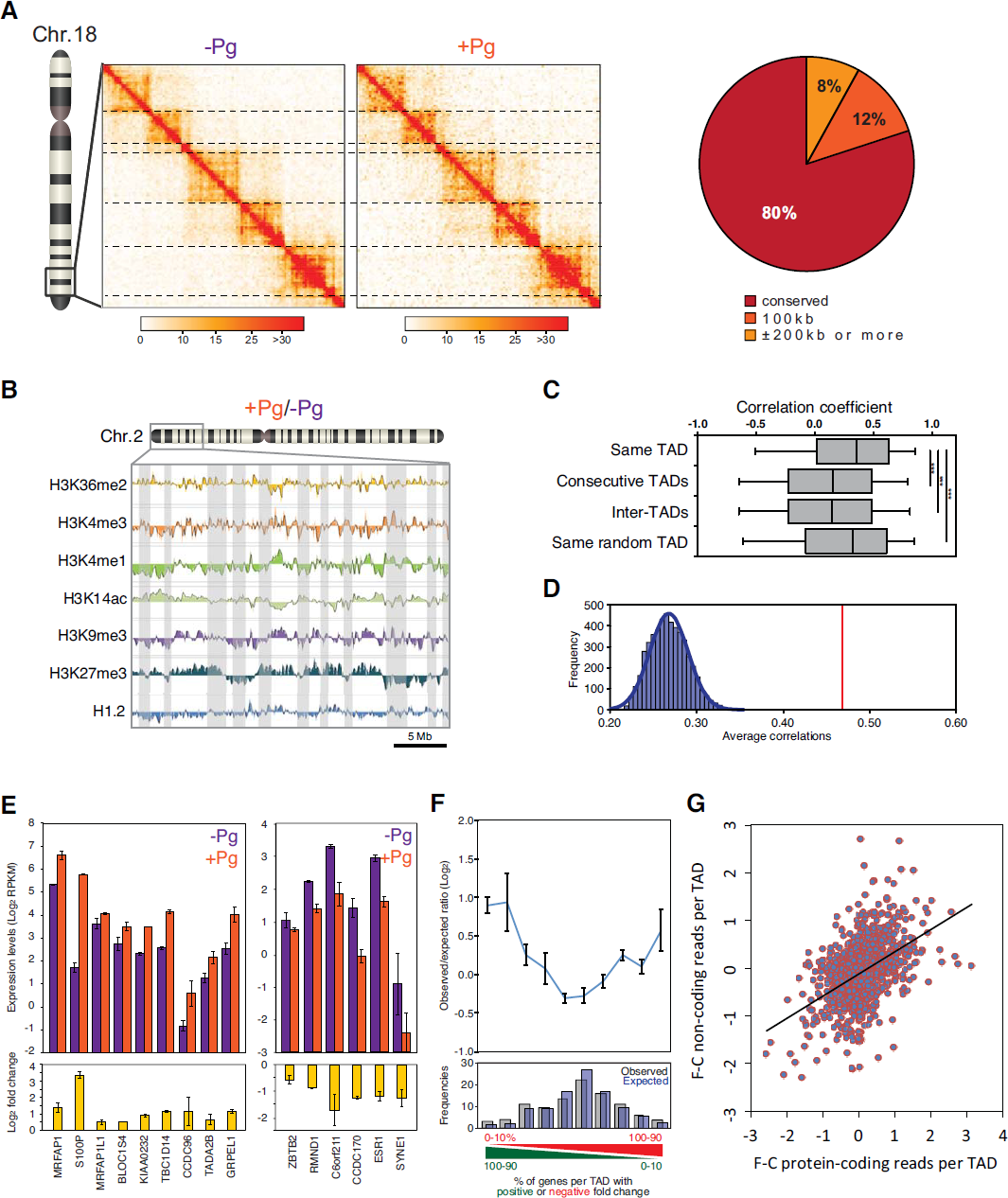
TADs chromatin and activity is coordinately modified upon hormone stimulation. **(A)** Chromatin interaction maps of a 10Mb region of the chromosome 18 before (-Pg) and after (+Pg) 1 hour of Pg treatment (Determined TAD borders are depicted by the horizontal dashed lines). The pie-chart shows the percentage of conserved borders between the -Pg and + Pg Hi-C datasets within 100Kb, or more than 100Kb. **(B)** Log_2_ Ratio of normalized ChIP-Seq signals + Pg/-Pg treatment over a 30 Mb region of the Chromosome 2. **(C)** Distributions of pair-wise correlation coefficients of Pg-induced chromatin profiles changes between 100Kb chromosomal fractions located within the same TAD, within consecutive or randomly picked TADs or in a similar randomly defined domain for the region depicted in B. (***) *P* < 0.001; Mann-Whitney test. **(D)** Genome-wide analysis of the pair-wise correlation between Pg-induced changes in epigenetic marks between 100 kb fractions belonging to a same TAD or to a same randomly positioned domain (see Supplementary Material). Histogram shows the distribution of genome-wide average correlation coefficients observed for 5,000 randomizations of TAD borders. The red bar (0.466) corresponds to the average correlation coefficients calculated for the real positions of the borders (*P* < 0.002). **(E)** Example of TADs with biased coordinated response to hormone, either activation (TAD 469 - left) or repression (TAD 821 - right). Upper panels show the expression levels of consecutive protein coding genes as determined by RNA-seq (average ± standard error of the mean) before (-Pg) or after 6 h of treatment with Pg (+Pg). Bottom panels show the ratio (log_2_) of expression of individual genes obtained after a 6 h treatment with Pg. **(F)** Top panel is the ratio of observed versus expected frequencies of TADs with distinct proportions of genes with positive or negative Pg-induced Fold Change FC > or < 1. Bottom histogram corresponds to the frequencies of TADs for the observed or randomized position of genes. **(G)** Pg-induced changes of expression of the pool of protein coding versus non-protein coding reads per TAD.

The observation above suggested that TADs could topologically constrain transient Pg-induced changes in gene expression. To explore this hypothesis, we performed two replicates of RNA-seq experiments in T47D treated or not with Pg for 6h (Supplementary Fig. 4A). We observed that some TADs have the majority of their genes either activated or repressed by the hormone (examples are shown in Fig. 2E). In addition to the thousands of genes significantly regulated by Pg (Ballare et al. 2013), we observed a general non-random distribution of divergent Pg-induced modifications of individual gene expression through the genome with stretches of genes which expression either slightly increases or decreases upon treatment. To generalize these observations, we calculated the percentage of genes with positive- or negative-fold changes after Pg treatment for 1,054 TADs with 4 or more protein coding genes. In this analysis, 178 TADs (∼17%) were found to harbor more than 80% of their genes changing expression in a similar direction (Fig. 2F), representing a significant enrichment as compared to expected random frequencies even after including robust randomizations of the gene positions that conserve an equivalent basal level of gene expression for each TAD (Fig. 2F ; *P* = 4.4×10^−7^ ; see Supplementary Material). In addition, independent analysis of the Pg-induced changes in RNA-seq reads for protein-coding and non-protein–coding regions located within the same TAD revealed that their changes of expression were significantly correlated (Fig. 2G ; Pearson R = 0.47 ; *P* < 0.001 ; see Supplementary Material and Supplementary Fig. 5). This indicates that genes with opposite response to Pg tend to be segregated in distinct TADs and that some TADs can be affected either by a global repression or stimulation of the expression of the genes they contain.

To characterize in a non-biased way TADs according to their activation or repression by Pg, we ranked all TADs with 4 or more protein coding genes according their global Pg-induced changes of expression and studied the top and bottom deciles (further mentioned as Pg-activated or -repressed TADs, respectively - see Supplementary Material) as compared to the remaining TADs (Fig. 3A and Supplementary Fig. 4a; see Supplementary Material). In these top and bottom TADs, the expression levels of most protein-coding genes (∼70%) were modified according to the global TAD response (Fig. 3B). These changes were irrespective of their basal expression levels (Supplementary Fig. 4B). In addition, the transcribed non-coding regions responded to Pg in the same direction as the protein-coding genes (Fig. 3C and Supplementary Fig. 5). We observed coherent activity changes already after 1 h of Pg treatment (Fig. 3D), whereas no correlated response to E2 was observed, suggesting that TADs could respond differently to distinct stimuli (Fig. 3E and Supplementary Fig. 4C). These results confirm that some TADs respond as a unit to hormone stimulus by global changes of their transcriptional activity.

**Figure 3:**
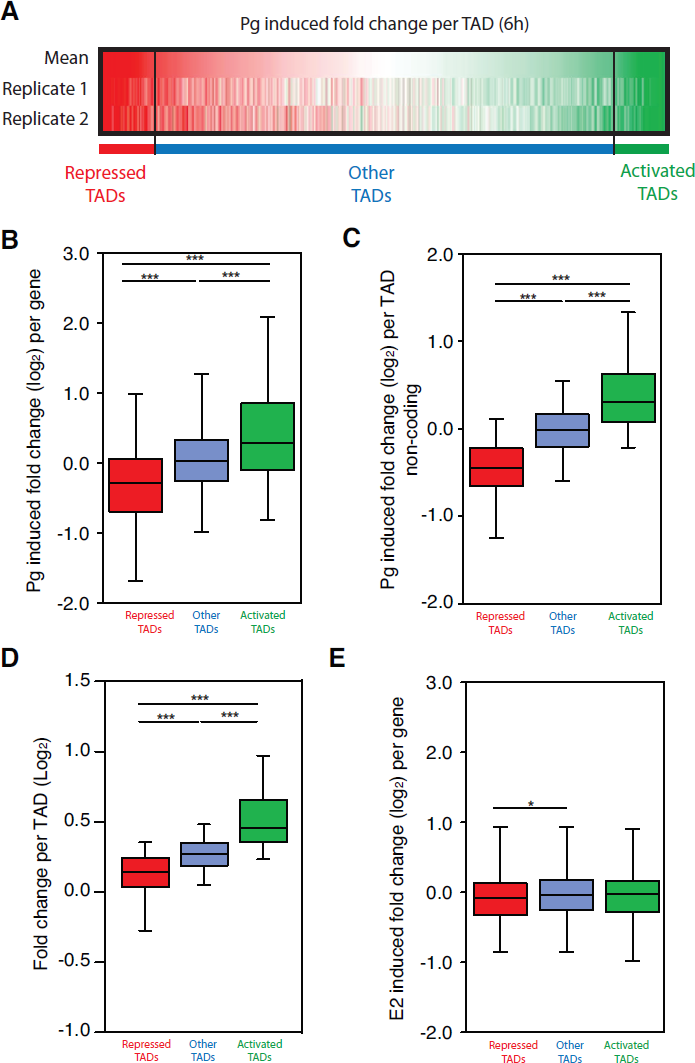
TADs respond as units to hormone in a stimulus-specific manner. **(A)** TADs ranked according their global changes of expression after a 6 h treatment with Pg. Repressed or Activated TADs are marked as the extreme deciles of the distribution (see Supplementary Material). **(B)** Distributions of Pg-induced fold changes of expression of protein coding genes in the different types of TADs after a 6 h treatment with Pg. **(C)** Distributions of Pg-induced fold changes of expression for regions that do not contain any annotated protein coding gene in the different types of TADs after a 6 h treatment with Pg. **(D)** Distributions of fold changes of TAD global expression induced by 1 h treatment with Pg for the three types of TADs. **(E)** Distributions of fold changes of TAD global expression induced by 6 h treatment with Estradiol (E2) for the three types of TADs. Boxplot whiskers correspond to 5^st^ and 95^th^ percentiles. (***),(**),(*) indicate P < 0.0001, 0.001 and 0.01, respectively (Mann-Whitney test).

### Pg induces structural reorganisation of TADs depending on the changes of activity of their resident genes

Next, we asked whether these coordinated transcription changes could be due to differential structural modifications of Pg-activated and -repressed TADs. Indeed, Pg treatment resulted in increased and decreased number or interactions within activated and repressed TADs, respectively (Fig. 4A). The correlation between changes in transcriptional activity and changes in interactions frequencies was observed independently in the two Hi-C datasets (*HindIII* and *NcoI*), supporting their Pg-dependence (Supplementary Fig. 6A, B).

**Figure 4:**
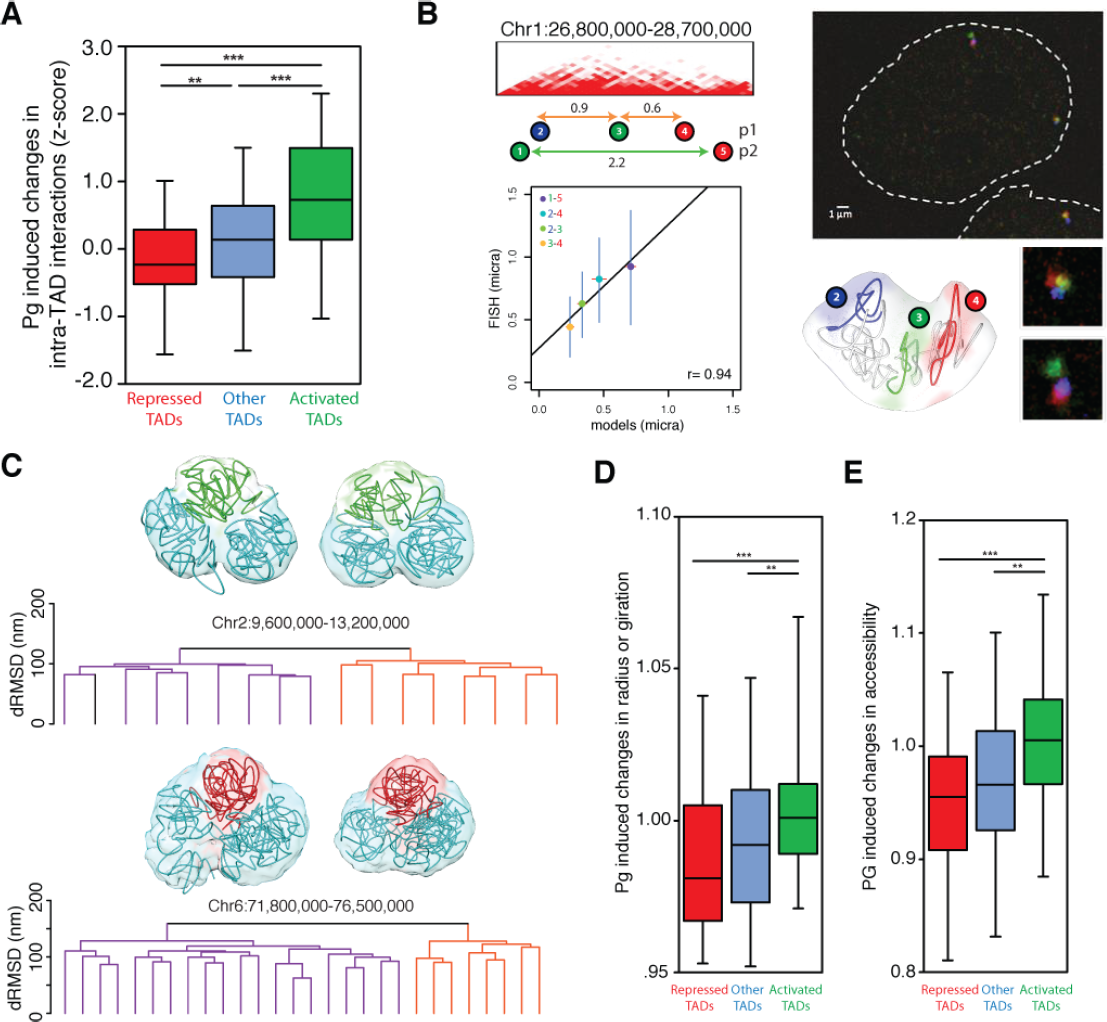
TADs undergo structural transitions upon hormone treatment. **(A)** Distributions of changes for intra-TAD interactions after 1 h treatment with Pg in the three types of TADs. **(B)** Two pools of BAC probes located within (p1) or beyond the borders (p2) of a diploid TAD of Chr.1 were used in 3D-FISH experiments (top right image show a representative result obtained with the p1; magnifications are shown below). Virtual 3D-FISH were performed on 3D models of the same region (bottom left panel). The mean and standard deviation of pair-wise inter-probes 3D distances obtained *in situ* (n = 60) were compared to the same distances obtained in the 3D models. **(C)** Three-dimensional models of activated TAD 170 (central green TAD top panel) and repressed TAD U767 (central red TAD bottom panel). Bottom graphs are dendrograms of the similarities between the resulting structural clusters of all models obtained for the TAD before (-Pg, purple) and after (+Pg, orange) treatment. **(D)** Distributions of Pg-induced changes of radius of gyration calculated from the models for the different types of TADs. **(E)** Distributions of Pg-induced accessibility changes calculated from the models for the different types of TADs.

To interpret these changes in terms of spatial structure-function relationship, we generated comprehensive three-dimensional (3D) models of 61 genomic regions of the T47D genome corresponding to 209 TADs, of which 30 and 43 were transcriptionally repressed and activated by Pg, respectively (Supplementary Table 3). Models were built based on our Hi-C datasets using the Integrative Modelling Platform (Russel et al. 2012) (IMP) as previously described (Bau and Marti-Renom 2012) (See Supplementary Material). We pooled the normalized Hi-C datasets obtained with *NcoI* and *HindIII* and used the interaction matrices at a resolution of 20 Kb, resulting in about 50 interacting particles per modeled TAD. To validate the modeling approach, we compared 3D models obtained for a region of chromosome 1 with the results of FISH experiments on the same region. We used five fluorescent probes located within the same TAD and/or near the borders (Fig. 4B). As previously described (Mateos-Langerak et al. 2009), the spatial distances measured *in situ* increased with the genomic distances (Supplementary Fig. 6C). This trend was also observed in the 3D models, albeit with smaller distances across the entire set of pairs of probes (Supplementary Fig. 6C). Despite the scale difference, the distances calculated from the 3D models correlated with the 3D FISH distances (Fig. 4B, R = 0.94) confirming that the accuracy of our modeling approach was appropriate to further analyze global structural properties of the modeled TADs.

We built 2,000 models for each of the 61 genomic regions and analysed the top 1,000 models (*i.e.* with the lowest IMP objective function). Hormone treatment modified the overall structure of TADs, such that the Pg repressed TAD models could not be structurally superimposed on the Pg activated TAD models (Fig. 4C). These results could give a structural base to the observations made on the Pg-induced changes of intra-TAD interactions (Fig. 4A). We first calculated the radius of gyration from our 3D models for each TAD before and after treatment. We observed that Pg-repressed TADs had a significant decrease in radius with respect to activated TADs (*P* < 0.002) (Fig. 4D). TADs with increased radius of gyration after Pg were likely to undergo structural decompaction, while TADs with decreased radius of gyration after Pg treatment were likely to adopt a more compact conformation (Fig. 4C). We hypothesized that Pg-induced increases and decreases in gyration radii of the TADs might affect their general accessibility. To independently test this hypothesis, we calculated the accessibility of single particles or whole TADs in our models (see Supplementary Material). Interestingly, we observed that model particles containing TSS sites were more accessible than the rest of the particles within the modeled TADs (Supplementary Fig. 6D; *P* = <0.001), suggesting a relationship between transcriptional activity and site accessibility within TADs and demonstrating that our approach captured functional parameters of the TADs organization. We therefore compared the Pg-induced accessibility changes for the 209 modeled TADs and observed that activated TADs had significantly increased accessibility with respect to non-affected or repressed TADs (*P* = 0.006 and *P* = 0.001, respectively - Fig. 4E). In summary, the 3D models indicate that TAD structures are modified in response to hormones, and that this is correlated with the changes in transcriptional activity.

### Structural transitions are associated with coordinated Pg-induced modifications of chromatin

The observed structural conversions may be linked to Pg-induced modifications of the structure and plasticity of chromatin within the responsive TADs (Vicent et al. 2008; Wright et al. 2012). To analyze this possibility, we compared the Pg-induced changes of chromatin components and epigenetic marks (See Supplementary Material) of activated and repressed TADs. Repressive marks accumulated in repressed TADs, whereas activating marks increased in hormone-activated TADs and decreased in repressed TADs (Fig. 5A). No coherent changes were observed in non-regulated TADs (Fig. 5A). These results correlate with the changes in accessibility from our 3D models and thus suggest that the Pg-induced chromatin changes participate in the structural transitions by modulating the flexibility of the chromatin fiber. For instance, depletion of H1 content has been associated with cooperative chromatin decondensation and may facilitate loop formation (Buttinelli et al. 1999; Routh et al. 2008; Diesinger et al. 2010). To assess whether the activated progesterone receptor (PR) drove these chromatin and structural modifications, we analyzed the distribution of PR binding sites (PRbs) as determined by ChIP-seq after 5, 30 and 60 minutes of Pg treatment (Fig. 5B). PRbs density was higher in activated TADs than in other TADs at any time point, in agreement with our previous gene-oriented studies (Ballare et al. 2013). At early time points, however, PRbs density was also higher in repressed TADs, suggesting that functional binding of PR could be the initial step leading to the structural rearrangements of TADs. Together our data suggest that Pg-repressed TADs are in a relatively open conformation prior to hormone treatment and become more compacted upon hormone addition, likely due to increases in H1, H2A and HP1 content. In contrast, Pg-activated TADs expand and de-compact after hormone treatment, likely due to depletion of H1 and H2A that increases chromatin fiber plasticity (Fig. 5C).

**Figure 5:**
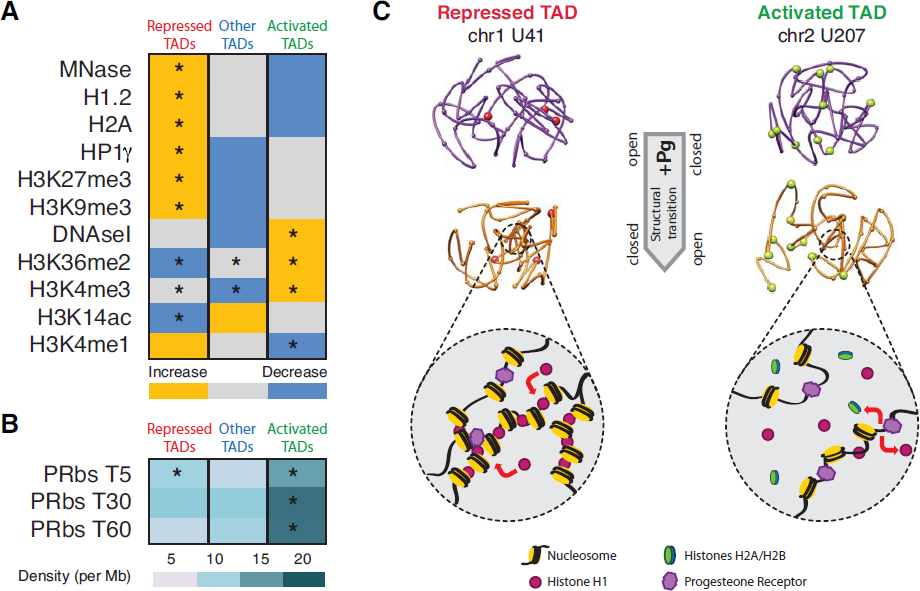
Pg-induced chromatin changes participate in the structural transitions. **(A)** Heatmap of the median ratio of normalised ChiP-seq signals with or without treatment for whole TADs classified as in Fig. 2 (relative Pg-induced decreases in blue and Pg-induced increases in orange). **(B)** Distributions of the density (per Mb) of Progesterone Receptor (PR) bound sites (PRbs) after 5, 30 and 60 min of treatment according the TADs categories. *: *P* < 0.05, Mann-Whitney test. **(C)** Model of Pg induced structural reorganization of responsive TADs: Repressed TADs become more compacted and their chromatin fibre gains core and linker histones, while activated TADs experience opposite changes.

## Discussion

Large-scale or genome-wide studies of chromosomes conformation by 5C, Hi-C or TCC, (Dostie et al. 2006; Lieberman-Aiden et al. 2009; Kalhor et al. 2012) revealed that the mammalian genome is organised in a hierarchy of structures, where chromatin globules (Bau et al. 2011) and Topologically Associating Domains (TADs) (Dixon et al. 2012; Nora et al. 2012) are integrated into chromatin compartments (Dekker et al. 2013). We found that in the T47D breast cancer cell line, this high level of genomic organization is conserved, including the partitioning of chromosomes into ∼1Mb-sized TADs. Since it has been proposed that TADs topologically separate chromatin domains with differential activity during differentiation (Nora et al. 2012), we hypothesized that it could also participate in transient gene regulation by constraining the nuclear environment of genes in terminally differentiated cells. The non-random organization of genes along eukaryotic chromosomes is well established and plays a role in the coordination of gene expression (Caron et al. 2001). Increasing evidences demonstrate that such coordination of expression might depend on looping-mediated interactions between regulatory elements and gene promoters, which could be located far away in the linear genome sequence (Bau et al. 2011; Shen et al. 2012; Thurman et al. 2012). We have used the response of a breast cancer cell line to a progesterone analogue (Pg) to explore a possible function of TADs in hormonal gene regulation. RNA-Seq analysis confirmed that Pg target genes are non-randomly located throughout the genome. Not only responsive genes are enriched in a limited number of TADs, but more generally genes differentially affected in response to Pg are frequently segregated in distinct TADs, which therefore could respond as units to the treatment. We also show that changes of gene expression are accompanied by structural transitions of the TADs, supporting the idea that TADs provide a structural base for local coordination of transient gene expression changes in differentiated cells.

We found that epigenetic states are more homogeneous within a TAD and that TAD borders frequently prevent the spreading of epigenetic features. Our analysis supports the concept that TADs represent homogeneous epigenetic domains with distinct permissiveness to transcription. In this context, cell-specific changes of the chromatin landscape within given TADs could delimit specific patterns of gene expression established mainly by the cell dotation of transcription factors. This would provide a mechanism to explain the frequent cell specific correlation of expression levels of neighboring genes (Caron et al. 2001).

We observed that the expression level of many genes is affected in a similar direction, although to a variable extent, within activated and repressed TADs. It is tempting to propose that such coordinated response depends on the high frequency of self-interactions within the limited space defined by the TAD borders. Stimuli activated enhancers or silencers would preferentially act on genes located within the same TAD and would influence coordinately their expression in a stochastic way. Such view is in line with observations made for the estrogen receptor using a ChIA-PET approach (Fullwood et al., 2009) and is supported by the fact that in activated TADs, PR binding sites with enhancer-like properties are enriched. Similarly, one could hypothesize on the existence of PR driven silencers within repressed TADs. These coordinated changes of expression might also reflect collateral ripples of modifications due to large-scale changes of the chromatin landscape initiated at the responsive sites since we show that Pg induced changes of chromatin states detectable over several 100Kb that are preferentially limited by TADs borders. Long-range remodeling of chromatin over Mb-sized domains has been previously reported in response to steroid hormones (Nye et al. 2002; Garcia-Becerra et al. 2010; Kocanova et al. 2010). Similarly, epigenetic modifications encompassing large domains of the genome have been described to participate in gene silencing and more recently in gene activation during the process of malignant transformation (Hon et al. 2012). Thus, TADs appear to limit specific changes in the combination of different chromatin markers, which might reflect local specific recruitment of histones modifying enzymes and chromatin remodeling activities. These findings, along with the epigenetic identity of the TADs, support the notion that the initial organization of TADs could act as topological constraint for changes in gene expression in response to external cues. Previous studies on the spatial organisation of glucocorticoid response in murine cells showed that the hormonal response was facilitated by a pre-organized conformation of the nucleus, particularly by cell-type-specific chromatin sites accessible to regulatory factors (Hakim et al. 2011). These nuclear hubs could reflect the organization in TADs, supporting an essential role of the 3D organization of the genome in the transcriptional response to hormone. The transient changes of gene expression in differentiated cells would be constrained by the chromatin state and 3D structure of TADs, which modulates access to genomic sequences for remodeling machineries as well as for specific and general transcription factors (Yadon et al. 2013). Such hypothesis is supported by the observations made recently that interactions between promoters and regulatory elements are established prior to treatment in the case of other type of transient stimuli (Jin et al. 2013).

Although similar boundaries demarcate TADs before and after Pg stimulation, our analysis demonstrates that the internal organization of TADs is modified upon hormonal stimulation. Indeed, TADs appear to re-structure internally after Pg induction and the magnitude of the structural changes depends on their chromatin state and correlates with the differential expression of the resident genes after hormone addition. Our 3D models suggest a direct functional influence of the conformation of TADs on the control of gene expression in response to external cues: TADs activated upon Pg treatment acquire a more open conformation that could facilitate the recruitment of general transcription machinery and/or the engagement of new regulatory contacts (i.e., increase of interactions). TADs that are transcriptionally repressed upon Pg treatment transition to a more compacted structure that might limit the accessibility to regulatory proteins. Further analysis with higher coverage and resolution is required to precisely map these changes in specific intra-TAD interactions between loci and their regulatory elements. However, it is probable that new chromatin loops are generated at the sub-TAD scale upon hormone stimulation that might require distinct combinations of chromatin organizing proteins (Phillips-Cremins et al. 2013). Overall, the large-scale Pg-induced structural changes were coherently correlated to the epigenetic modifications of the chromatin landscape. Therefore, our results indicate for the first time the existence of a link between dynamic structural changes, epigenetic markers and transcriptional regulation upon transient stimuli in differentiated cells.

Structural changes in Mb-size domains of the genome would intuitively require changes of the physical properties and plasticity of the chromatin fiber. Our previous works demonstrated that transcription regulation by Pg required specialized chromatin remodelling for both activation and inhibition of genes (Vicent et al. 2011; Wright et al. 2012). We show that Pg induced profound changes on the chromatin structure exemplified by a modification of the MNase sensitivity and content in core histone H2A as well as in the linker histone H1 (Ballare et al., 2013). Although the molecular mechanisms coordinating these Pg induced chromatin changes within TAD remain to be fully elucidated, our data suggest that the displacement of H1.2, which has been previously associated to cooperative de-condensation of chromatin (Buttinelli et al. 1999; Routh et al. 2008), could make the chromatin fiber thinner and more flexible within activated TADs. This would increase its capacity for exploring a larger territory in response to Pg, leading to the observed increase in chromatin interactions and to the structural expansion we observed in our models. In repressed TADs we observed the opposite behavior, increase in histone H1.2 content upon Pg treatment, which could lead to a thickening of the chromatin fiber and consequently to lower flexibility, decrease interactions and compaction of the TAD, limiting their general accessibility to regulatory proteins.

In summary, our analysis suggests that specialized TADs respond coordinately upon transient hormonal stimulation. Such coordinated response would depend on favoring self-interactions inside the limited space of TAD, facilitating a rapid response by locally reorganizing its structure. Importantly, an opposite structural reorganization is observed within activated and repressed TADs depending on the changes of activity of the resident genes. Hormone regulated differential loading/recruitment of histone H2A and linker histone H1.2 into the chromatin fiber offers a plausible molecular mechanism for regulating access to regulatory elements for transcription factors and other chromatin interacting protein complexes. We propose that TADs could be considered as units of integration of the numerous pathways that are regulated upon environmental changes. Such model would be of particular importance during the process of malignant transformation, when changes of the chromatin state or nuclear position of given TADs could have direct influences on the expression of its entire gene set. Additional experiments will be needed to elucidate whether similar events occur for other types of external stimuli, and whether inter-TAD interactions and nuclear localization modulate the response of individual TADs.

## Methods

### Cell culture and hormonal stimulation

All experiments (including Chip-seq, Hi-C and other high-throughput analyses described in this study) were performed in the following conditions of culture and treatments: T47D cells were grown 48 h in RPMI supplemented with 10% charcoal-treated FBS and synchronized in G0/G1 by 16 h serum starvation. The Pg analogue R5020 (at 10^−8^ M) was added to the medium, and cells were harvested after the times indicated.

### Epigenetic data collection and analysis

MNAse-seq, DNAse I-seq and ChiP-seq experiments (for H3K4me1, H3K4me3, H2A, H4, RNA polymerase II, progesterone receptor, H3K9me3, HP1γ) in T47D cells were described previously (Ballare et al. 2013; Vicent et al. 2013). ChiP-seq experiments for CTCF, H3K36me2, H3K27me3, H3K14ac and H1.2 were performed in similar conditions using the following antibodies: 07-729 (Millipore), 07-369 (Millipore) 39155 (Active Motif), 07-353 (Millipore) and ab4086 (Abcam), respectively. Reads were processed by aligning to the reference human genome (GRCh37/hg19). MNase-seq, DNase I-seq and ChIP-seq signals normalised for sequencing depth were summed according to different windows sizes (ranging from 100 Kb to whole TADs). Summed reads were divided by the corresponding signal obtained for an input DNA to determine the normalised signal over input enrichment or depletion. Similarly, progestin-induced enrichment or depletion of mark content was determined for 100 Kb bins or for the whole TAD as the ratio of sequencing-depth normalised read counts before and after treatment.

### Generation of contact matrices and integrative modelling of spatial contacts

Hi-C libraries were generated from T47D cells treated or not with R5020 for 60 min according the previously published Hi-C protocol with minor adaptations (Lieberman-Aiden et al. 2009). Hi-C libraries were generated independently in both conditions using *HindIII* and *NcoI* restriction enzymes. Hi-C libraries were controlled for quality and sequenced on an Illumina Hiseq2000 sequencer. Illumina Hi-seq paired-end reads were processed by aligning to the reference human genome (GRCh37/hg19) using BWA. Datasets normalized for experimental biases and sequencing depth (Lieberman-Aiden et al. 2009) were used to generate contact matrices at 20, 40 and 100 Kb as well as at 1 Mb resolutions.

### Classification of TADs according their transcriptional response to Pg

To classify TADs according their hormone response, we calculated the average ratio of the number of normalised RNA-seq reads obtained after and before hormone treatment in two RNA-seq replicates. TADs containing more than three protein coding genes were conserved for further analysis and ranked according the average ratio described above. The top and bottom 10% were classified as “activated” and “repressed” TADs, respectively. To independently test the homogeneous transcriptional response within TADs, we calculated the same ratio using the RNA-seq reads corresponding to either protein-coding genes or non-coding transcripts (about 10% of the reads per TAD in average).

### 3D modelling of genomic domains based on Hi-C data

Hi-C produces two-dimensional matrices that represent the frequency of interactions between loci along the genome. To transform such data into a 3D conformation of higher-order chromatin folding, we used the Integrative Modeling Platform (IMP, http://www.integrativemodeling.org) (Russel et al. 2012). Structure determination by IMP can be seen as an iterative series of four main steps: generating experimental data, translating the data into spatial restraints, constructing an ensemble of structures that satisfy these restraints and analysing the ensemble to produce the final structure. All 3D models were built as previously described (Bau and Marti-Renom 2012).

See Supplementary Material for further details and additional methods and procedures.

## Acknowledgments

We thank the CRG Ultra-sequencing and Advanced Light Microscopy Facilities for technical support, and all members of the Chromatin and Gene Expression and Structural Genomic groups for helpful discussions. We acknowledge Juan Valcarcel for his helpful comments on the manuscript. We also thank Marta Morell and the Genetic Causes of Disease group for providing the BAC clones used in this study. We acknowledge financial support from the Spanish MINECO (BFU2010-19310/BMC) and the Human Frontiers Science Program (RGP0044/2011) to M.A.M.-R. This work was also supported by grants from the Spanish government (BMC 2003-02902 and 2010-15313; CSD2006-00049) and the Catalan government (AGAUR) to M.B.

### Author Contributions

F.L.D., G.F., M.A.M-R. and M.B. designed the studies; F.L.D. performed Hi-C, FISH and RNA-Seq experiments; G.P.V., R.H.G.W and C.B. performed ChIP-Seq experiments. F.L.D. carried out the data analysis with contributions of A.P., D.S., G.C and G.F.; D.B. and M.A.M-R carried out the 3D modelling and analysis; F.S. and G.F. designed the TAD boundary algorithm; all authors discussed the results and F.L.D., M.A.M-R. and M.B. wrote the manuscript.

